# Theoretical origins of extreme cell differentiation irreversibility: a dynamic model of organismal size effects

**DOI:** 10.1101/2025.04.20.649670

**Authors:** Yuanxiao Gao, Xueyan Zhao, Caixia Li

## Abstract

Cell differentiation is a pivotal evolutionary transition underlying multicellular complexity. Irreversible differentiation – a process in which cells permanently lose plasticity to become specialized types – is hypothesized to emerge predominantly in large, cell-rich organisms. However, the causal relationship between organismal size and the evolutionary origins of this extreme developmental commitment remains unclear. By integrating developmental dynamics into a mathematical model, we demonstrate that extreme irreversible differentiation (where all cell types exhibit irreversibility) most robustly evolves in intermediately sized multicellularity. This reveals a non-monotonic dependence of cell fate determinism on an organismal scale. Specifically, increasing organismal size initially promotes extreme irreversible differentiation by enhancing the selective advantages of cellular division of labor. Beyond a critical size threshold, however, further expansion destabilizes this extreme irreversibility due to its decreasing differentiation advantages compared with reversible differentiation. Counterintuitively, smaller organisms establish cellular irreversibility at earlier developmental stages rather than larger organisms. Our framework identifies an evolutionary double-edged sword: while body size expansion initially facilitates extreme differentiation irreversibility, it ultimately leads to its collapse. These results establish quantitative links between organismal dimensions and cell fate determinism, explaining the size-dependent evolution of developmental strategies in simple multicellular systems.

## 1 Introduction

Large body size confers multifaceted adaptive advantages across life’s organizational hierarchy, spanning microbial colonies to complex metazoans [1–7]. A central advantage of attaining large size in multicellular systems is the capacity for cellular differentiation – defining innovation enabling biological complexity through specialized cell type generation [8– 10]. This developmental mechanism drives hierarchical organization by partitioning cellular functions, thereby escalating organismal complexity [11]. While unicellular organisms exhibit limited capacity for temporal differentiation producing phenotypically distinct states for stage-specific tasks [12, 13], they lack the spatial differentiation machinery required for coordinated developmental regulation [14]. For instance, *Naegleria gruberi* demonstrates transient phenotypic switching between flagellated and non-flagellated forms [13]. In contrast, multicellular organisms leverage their numerical superiority to establish stable cell type specialization through developmental commitment. Studies confirm that organismal size positively correlates with cell type diversity, serving as a key metric of biological complexity [8, 11, 15, 16].

Irreversible differentiation represents a terminal commitment process wherein cells progressively lose phenotypic plasticity to generate specialized cell types with fixed functional roles [4, 9, 17–21]. This is exemplified in filamentous cyanobacteria, where vegetative cells irreversibly differentiate into nitrogen-fixing heterocysts triggered by nitrogen deprivation [22]. The evolutionary stability of such irreversible specialization hinges on two prerequisites: (i) functional integration within multicellular architectures, and (ii) compensatory coordination between distinct cell lineages. Theoretically, larger organisms accommodate more elaborate irreversible differentiation networks through spatial partitioning and metabolic channeling. Comparative analyses of volvocine algae demonstrate a size-dependent continuum of differentiation strategies: large colonies (*>*500 cells) evolve discrete germ-soma differentiation through terminal commitment and small colonies (*<*50 cells) maintain undifferentiated cells, revealing threshold-driven constraints on the evolution of differentiation irreversibility [4]. This size-dependent continuum suggests developmental constraints shape differentiation evolvability, yet the evolutionary mechanisms linking size thresholds to differentiation irreversibility remain unresolved.

The evolutionary emergence of cellular differentiation has been widely modeled [6, 19, 23–30] and experimentally validated [2], yet the mechanistic links between organismal size and differentiation irreversibility remain unresolved. While organismal size has been recognized as a potential modulator of differentiation patterns [6, 28, 31], existing studies present conflicting insights: topological constraints nullify size effects in ring-structured colonies but amplify them in tree-like structured organisms [28], whereas coordination costs favor specialized small groups [31] and cancer risks shape differentiation strategies in larger organisms [6]. Parallel work on cell differentiation mechanisms emphasizes cellular interactions [19, 32] but neglects size-dependent selection pressures. This disconnect persists despite evidence that terminal differentiation irreversibility scales with cellular type complexity [32], highlighting the need to integrate organism size with evolutionary cell differentiation models.

In this study, building upon previously established theoretical models [19, 21], we systematically investigate organismal size effects on the evolution of irreversible cell differentiation. Our computational framework defines organismal size as the maximal number of cell divisions allowed during development, a parameter that is explicitly defined in the model. Cellular differentiation dynamics are modeled through distinct division patterns emerging during organismal growth. We specifically focus on extreme irreversible differentiation (*EID*), which is characterized by progressive loss of differentiation potential across all cellular lineages. These *EID* patterns compete with alternative differentiation strategies through natural selection, which acts to maximize organismal fitness, quantified here as the expected reproduction rate throughout the life cycle. Our investigation progresses through three principal axes: First, we establish how organismal size shapes the evolution of *EID*. Second, we delineate size-dependent variations in the temporal progression of cellular irreversibility during developmental stages. Finally, we extrapolate these dynamics to evolutionary scenarios involving organisms approaching biological size limits.

## 2 Model

### Life cycle of organism

To analyze the size-dependent evolution of differentiation patterns, we extend a multicellular model [19, 21] with organismal size variation. The framework integrates volvocine algae observations [4] showing size-differentiation correlations: undifferentiated (small), partially specialized (medium), and fully specialized (large) species. Building upon the division of labor hypothesis [13, 23, 33], we postulate two transitional cell states: reproductive (*R*) and somatic (*S*) lineages, with functional specialization aligned with previous characterizations [19, 21, 23, 31, 33]. These cell types exist in transitional states enabling bidirectional interconversion, contrasting with terminally differentiated cells locked in determined developmental trajectories. Organisms initiate from a founder *R* cell (Fig. 1A), proliferate through *n* synchronous cell divisions until reaching size maturity size with 2^*n*^ cells. Upon maturation, *R* cells undergo morphogenesis to establish new progeny, while *S* cells execute cell death. This reproductive cycle iterates through successive generations.

**Figure 1:**
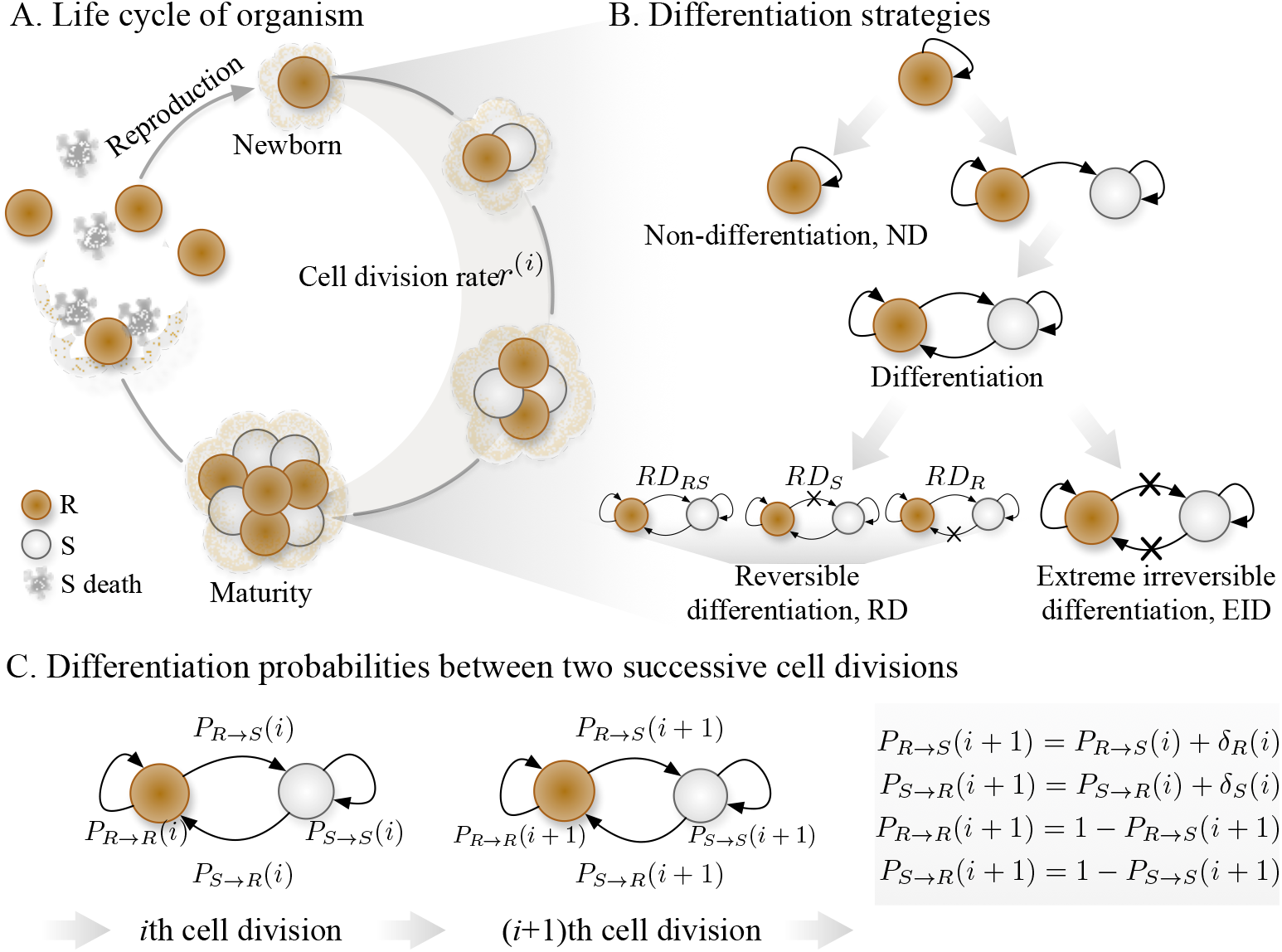
Illustration of an organism’s life cycle, differentiation strategies, and differentiation probabilities between two cell divisions. The life cycle of each organism (**A**) initiates with a single *R* cell, proceeding through synchronous developmental cell divisions before reproductive maturity. At reproduction, all *S* cells perish, leaving only *R* cells as offspring. Differentiation strategies (**B**) are defined by cell-type plasticity during development: non-differentiation (*ND*) occurs if *R* cells exclusively generate *R* cells, while reversible strategies (*RD*) involve bidirectional or unidirectional transdifferentiation. Specifically, *RD*_*RS*_ permits mutual conversion between *R* and *S* cells at the *n*th division, whereas *RD*_*R*_ and *RD*_*S*_ restrict reversibility to *R*-to-*S* or *S*-to-*R* transitions, respectively. In contrast, extreme irreversible differentiation (*EID*) enforces lineage commitment, where each cell type exclusively self-replicates. Differentiation probabilities (**C**) are quantified as *P*_*R*→*S*_(*i*) and *P*_*S*→*R*_(*i*), denoting the likelihood of *R*-to-*S* or *S*-to-*R* conversion at the *i*th division. Between successive divisions (*i*) and (*i* + 1), these probabilities may diverge by increments *δ*_*R*_(*i*) and *δ*_*S*_(*i*), such that *P*_*R*→*S*_(*i* + 1) = *P*_*R*→*S*_(*i*) + *δ*_*R*_(*i*) and analogously for *P*_*S*→*R*_(*i*).

### Differentiation strategy

Differentiation strategies are governed by temporal sequences of cellular transition probabilities throughout organismal development (Fig. 1B), with stochasticity quantified via differential increments *δ* between successive divisions (Fig. 1C). The Markovian dynamics follow: *P*_*R*→*S*_(*i*+1) = *P*_*R*→*S*_(*i*)+*δ*_*R*_(*i*), where *P*_*R*→*S*_(*i*) denotes the *R*– to–*S* transition probability at the *i*th cell division. Analogous formula defines *S*–to–*R* transitions: *P*_*S*→*R*_(*i*+1) = *P*_*S*→*R*_(*i*)+*δ*_*S*_(*i*). These probability trajectories 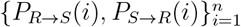 formally define differentiation strategies [21]. Differentiation strategies are classified through three operational criteria: (1) Non-differentiation (ND) occurs when all *R*–to–*S* transition probabilities remain null *P*_*R*→*S*_(*i*) = 0 for *i* = 1, …, *n*); (2) Extreme irreversible differentiation (EID) requires differentiation events with terminal state fixation (*P*_*R*→*S*_(*n*) = *P*_*S*→*R*_(*n*) = 0); (3) Reversible differentiation (RD) encompasses all other scenarios, which are further partitioned into three subclasses based on terminal state distributions: bidirectional reversible (*RD*_*RS*_), somatic revisable (*RD*_*S*_), and reproductive reversible (*RD*_*R*_) as illustrated in Fig. 1B. Of particular note, the *RD*_*R*_ is previously characterized as irreversible somatic differentiation due to the irreversibility in somatic lineage [21]. Notably, the stochastic perturbation parameters (*δ*_*R*_(*i*), *δ*_*S*_(*i*)) act as phase space controllers – their nullification (*δ*_*R*_(*i*) = *δ*_*S*_(*i*) = 0) reduces the system to stage-independent differentiation dynamics consistent with prior deterministic frameworks [19, 21].

### Fitness

The evolutionary optimization of differentiation strategies is formalized through fitness maximization dynamics, where organismal fitness is operationalized as the expected reproduction rate *λ*(*n*) – quantified by the exponential growth rate of reproductive (*R*) cells [19, 21]. Following established frameworks [21], we define stage-dependent growth kinetics through the benefit-cost ratio:

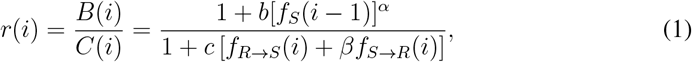

where *f*_*x*_(*i* − 1) denotes the proportion of *x*-type cells after the (*i* − 1)*th* division, and *f*_*x*→*y*_(*i*) = *f*_*x*_(*i* − 1)*P*_*x*→*y*_(*i*) represents differentiation flux between types *x, y* ∈ {*R, S*}. Benefit composition (*B*(*i*)) and cost composition (*C*(*i*)) measure the contribution of *S* cells and the differentiation costs to the cell division rate, respectively. *α* and *β* are weight factors of benefit and cost. The recursive cellular dynamics are governed by:

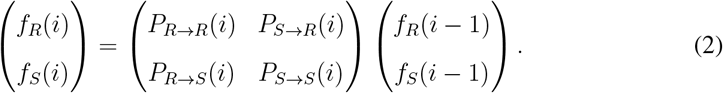

The developmental timeline is characterized by the cumulative maturation duration:

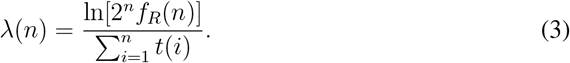

where 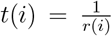 is the expected waiting time between the (*i* − 1)th division and the *i*th division [34]. Initial conditions are defined by *f*_*R*_(0) = 1 and *f*_*S*_(0) = 0, reflecting the *R*-cell progenitor paradigm. The non-differentiation (ND) strategy establishes the biological baseline with *λ*_*ND*_(*n*) = ln 2 through developmental invariance (*f*_*R*_(*i*) ≡ 1). Optimal strategy identification proceeds through comparative analysis of *λ*(*n*) across (*b, c, α, β*) parameter space, with particular focus on phase transitions in differentiation commitment dynamics (Appendix A.1).

## 3 Results

### 3.1 Extreme Irreversible Differentiation Reaches Evolutionary Optimum at Intermediate Sized Organisms

Our analysis reveals a non-monotonic relationship between organismal size and evolutionary selection pressures, with intermediate-sized organisms optimally promoting extreme irreversible differentiation (*EID*) emergence (Fig. 2A). The evolutionary accessibility of extreme irreversible differentiation (*EID*) is fundamentally constrained by its terminal differentiation requirement (*P*_*R*→*S*_(*n*) = *P*_*S*→*R*_(*n*) = 0), which biologically restricts its emergence to organisms with *n <* 2 developmental stages. Then, we found that *EID* evolved in the parameter regime where *b* and *c* are comparable (Fig. 2A). To elucidate this phenomenon, we first examine the optimal strategies under varying *b* and *c* via reproductive rate. The reproductive success metric *λ*(*n*), defined in Eq. (13), depends critically on the interplay between cellular division rates *r*(*i*) and offspring yield 2^*n*^*f*_*R*_(*n*).

**Figure 2:**
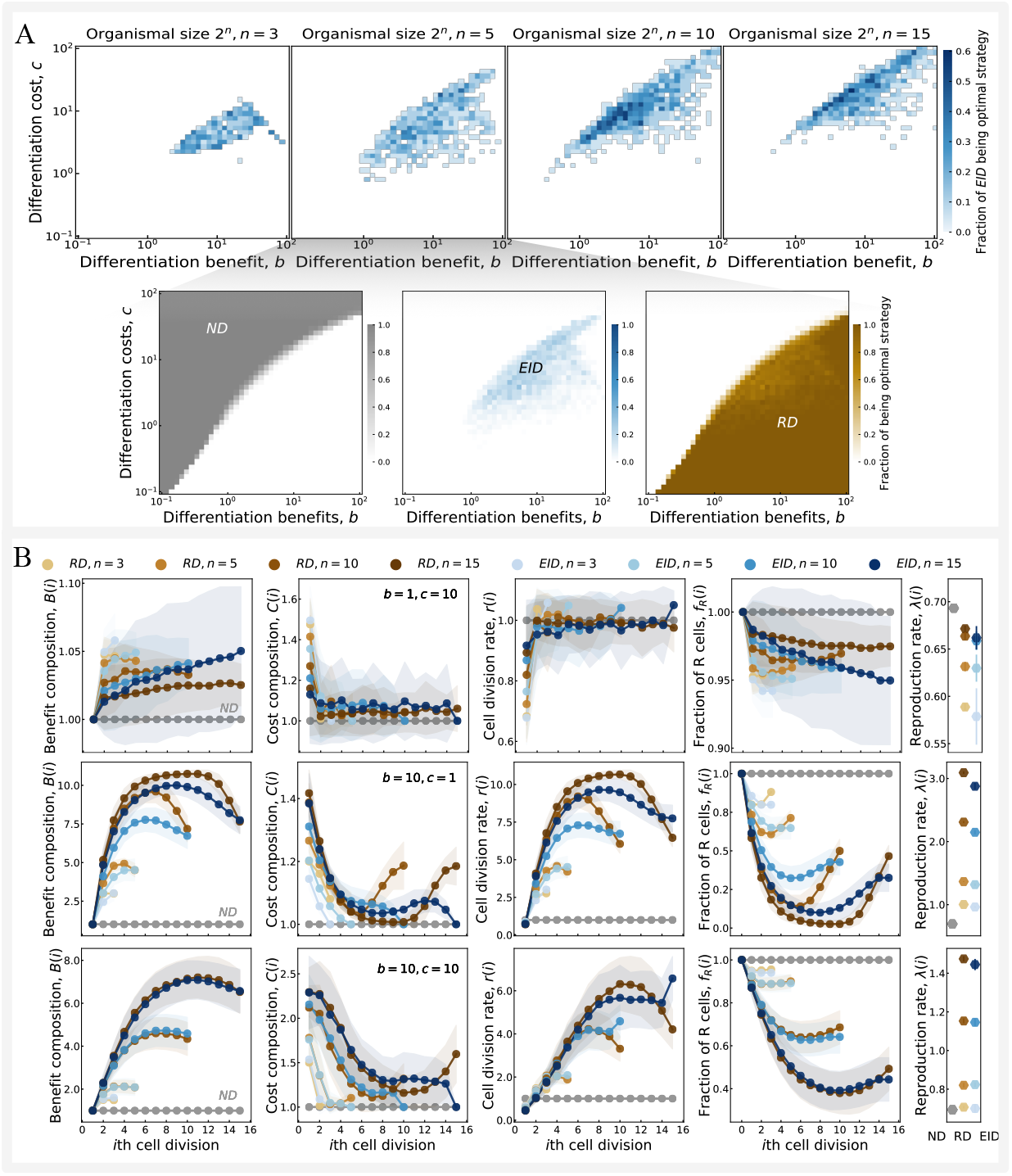
Organismal size changes the evolutionary likelihood of extreme irreversible differentiation. A. The fraction of each strategy (*ND*/*RD*/*EID*) achieving optimality under differentiation benefits *b* and costs *c* varies systematically with organismal size. **B**. Characteristics of optimal reversible differentiation (*RD*) and extreme irreversible differentiation (*EID*) (together with *ND*) reveal size-dependent patterns: differentiation benefit composition *B*(*i*), cost composition *C*(*i*), cell division rates *r*(*i*), *R*-cell proportion *f*_*R*_(*i*), and reproduction rate *λ*(*n*) exhibit distinct scaling relationships with organismal size. Data points (dots) and shaded areas represent the mean and the standard deviation for each parameter. Parameter specifications for panel **A** included 20 independent replicates for *n* = {3, 5, 10, 15} in the upper subpanel and 100 replicates for *n* = 5 per pixel in the lower subpanel. Developmental trajectories in panel **B** were derived from 20 replicates per optimal strategy. For more detail of the calculation of *λ*(*n*), see appendix A.2.

Under high cost-to-benefit ratios (*c* ≫ *b*), both reversible differentiation (*RD* and *EID*) strategies incur net fitness losses through excessive somatic commitment. Optimal solutions in this regime minimize *S*-cell production (1 − *f*_*R*_(*i*)) to reduce costs, and thus to maintain division rates (upper panels, Fig. 2B). Consequently, non-differentiation (*ND*) emerges as the dominant strategy through its avoidance of differentiation costs. Conversely, when *c* ≫ *b, RD* strategies maximize fitness by strategically accumulating *S*-cells during mid-development to enhance division rates, while preserving sufficient *R*-cells (*f*_*R*_(*n*)) for reproduction (middle panels, Fig. 2B). *EID* occupies the evolutionary interface between these regimes, exhibiting hybrid dynamics: early-stage differentiation resembling *RD* for benefit acquisition, followed by terminal *ND*-like reproductive conservation (Fig. 2B bottom panels). This strategic duality enables *EID* to outperform *ND* and *RD* within specific *b*-*c* phase boundaries, consistent with prior findings [21].

While large organisms theoretically permit *EID* to be optimal for maximal reproduction rates, our analysis reveals a paradoxical evolutionary outcome: reversible differentiation (*RD*) maintains dominance in large developmental organisms (*n >* 10) through its unconstrained probability space, whereas *EID*’s terminal irreversibility constraints as organismal size increases. This result stems from the dual dependence of fitness *λ*(*n*) on both cell division rates *r*(*i*) and offspring number 2^*n*^*f*_*R*_(*n*). Crucially, *RD*’s reproduction rate grows more substantially through strategic late-stage *R*-cell preservation to outweigh *EID*’s. This evolutionary landscape confirms the Goldilocks principle for cellular commitment strategies– maximal *EID* optimization occurs at intermediate biological scales. Notably, large organisms (*n* ≥ 10) exhibit expanded *EID* viability domains under elevated differentiation costs (*c* ≥ 10^2^), as shown in Fig. 2A. Although *r*(*i*) decreases under higher *c* values, the extended developmental procedure required by larger *n* leads to greater offspring numbers (2^*n*^*f*_*R*_(*n*)). This reproductive advantage ultimately outweighs the reduction in *r*(*i*) for *λ*(*n*) when compared with *ND* strategies. The resultant size-dependent advantage displaces *ND* strategies at high differentiation costs, as demonstrated in Fig. 2A.

### 3.2 Organismal size governs the temporal establishment of developmental irreversibility

Our analysis reveals that organismal size governs cellular irreversibility timing during development (Fig. 3A). Cellular differentiation states emerge through dynamic interactions between differentiation benefits, costs, and organismal scale parameters. Notably, size exerts dominant control over irreversibility patterns in larger organisms (*n* ≥ 15), resulting in delayed cell irreversibility for both cell types (Fig. 3A). Under small organism sizes (*n* = 3), differentiation benefits and costs have limited roles in the cell differentiation stages of both types. While as organism size increases (*n* = 5, 10), with low differentiation costs (*c <* 10), we observe distinct temporal differentiation patterns: *R* cells exhibit late-stage irreversibility while *S* cells differentiate earlier. This strategic prioritization of *S*-cell production maximizes reproductive success through accelerated division rates under low differentiation costs. The optimal differentiation strategy consequently features high early-stage *P*_*R*→*S*_(*i*) values with suppressed *P*_*S*→*R*_(*i*) (Fig. 3B, left panel), reflecting developmental constraints imposed by initial *R*-cell setting. Contrastingly, elevated differentiation costs (*c* ≥ 10) select against *S*-cell proliferation, favoring strategies with reduced differentiation probabilities throughout development (Fig. 3B, right panel).

**Figure 3:**
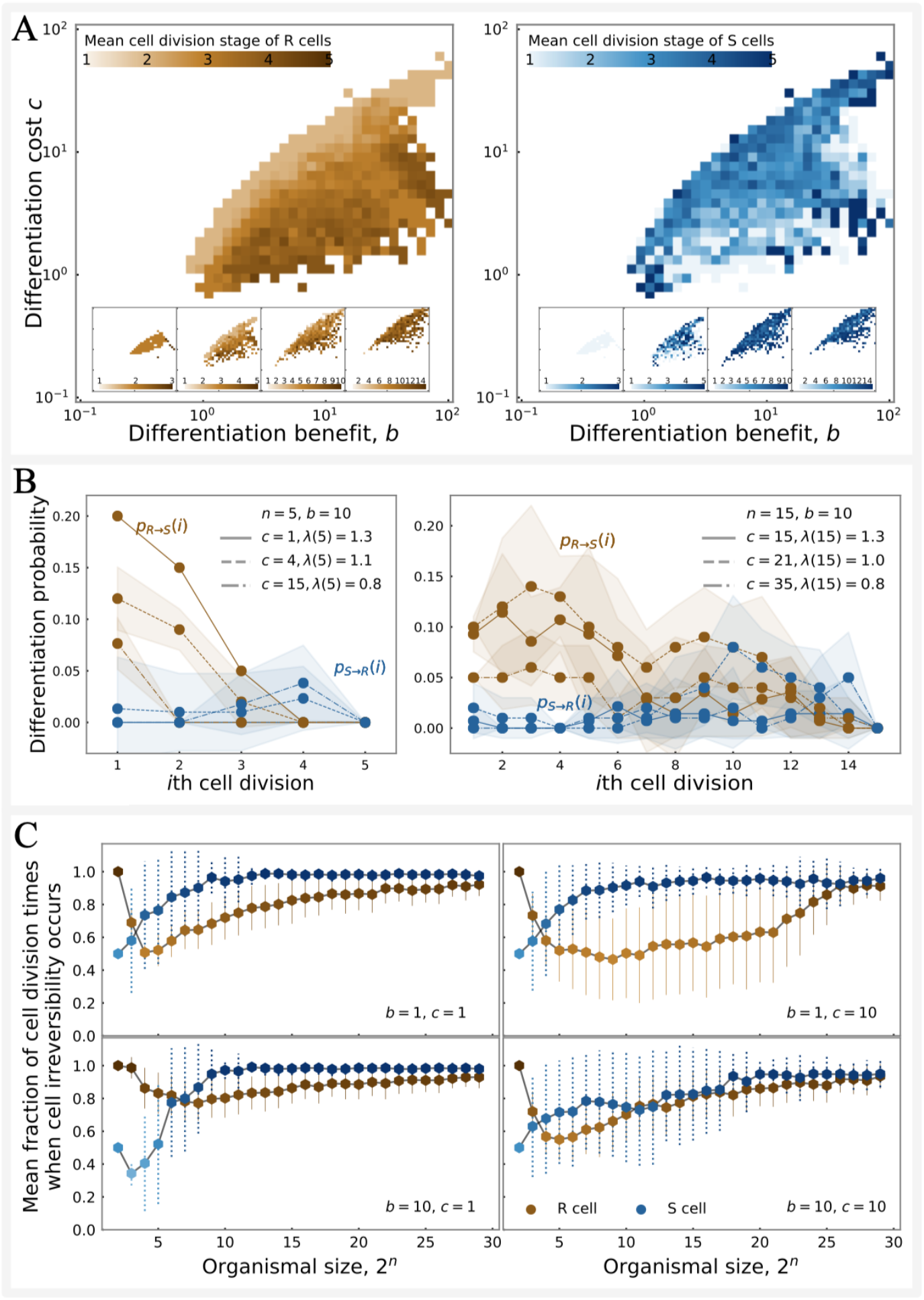
Developmental-stage timing of extreme irreversible differentiation. The mean cell division stages of *R* and *S* cells when *EID* constitutes the evolutionarily optimal strategy are shown in **A**, with colors indicating normalized division stages (scaled by organismal size *n*) at which irreversibility emerges. Insets display mean division stages across organismal sizes ranging from 2^3^ to 2^15^. Differentiation probabilities and reproductive rates of optimal *EID* strategies under varying benefit-cost ratios and organismal sizes are quantified in **B**, where brown and blue circles denote *P*_*R*→*S*_ and *P*_*S*→*R*_ probabilities, respectively. **C** presents the mean developmental-stage distribution of extreme irreversible differentiation events among the top 100 *EID* strategies (ranked by reproduction rate, though not necessarily globally optimal), with circles marking averaged differentiation stages of cell irreversibility and error bars representing standard deviations. Parameters: initial sampling size *M* = 1000, strategy sampling *SAM* = 100, with 100 replicates per pixel in **A** and 20 in **B/C**; full numerical procedures are detailed in Appendix A.2.

However, increasing organismal size delays the establishment of cellular irreversibility to later cell division stages (Fig. 3A, B). While both cell types achieve irreversibility during terminal divisions, optimal *EID* strategies exhibit relatively high *P*_*R*→*S*_ probabilities in early divisions and low probabilities in late divisions (Fig. 3B). This pattern demonstrates conserved differentiation probability dynamics - preferential *S*-cell production by *R* cells early in development, reversing in later stages. Larger organisms with extended division sequences leverage flexible differentiation to optimize cellular composition, achieving comparable reproductive rates to smaller organisms despite elevated differentiation costs (Fig. 3B). For instance, *λ*(15) has the same reproduction rate as *λ*(5) even under a higher differentiation cost.

To systematically examine organismal size effects on cellular irreversible differentiation, we analyzed division stages establishing irreversibility in *EID* strategies exclusively. Our results reveal that maximal-fitness *EID* strategies universally exhibit late-stage irreversibility establishment (Fig. 3C), though specific irreversible stage exhibits parameter-dependent shifts across *b* and *c* values. Under *c*-high/*b*-low parameter regimes, the division stages of irreversibility show high divergence between cell types. High variations of irreversible differentiation stages reflect multiple *EID* configurations achieving equivalent fitness maxima. Notably, larger organisms also demonstrate reduced variation in the irreversibility patterning establishment. These findings collectively demonstrate that increased organismal scale favors selection for *EID* strategies with delayed cellular irreversible differentiation.

### 3.3 Body size limits the evolution of extreme irreversible specialization

Theoretical investigation of strategy competition as *n* approaches infinity reveals critical insights into evolutionary dynamics. Previous simulation studies have shown that “irreversible somatic differentiation” gains selective advantage with increasing organismal size when differentiation probabilities remain constant (*δ*_*R*_(*i*) = *δ*_*S*_(*i*) = 0) [19]. Within this *δ* = 0 framework, three distinct strategy categories emerge: *ND* (no differentiation), “irreversible somatic differentiation” (defined by *P*_*S*→*R*_ = 0 with *P*_*R*→*S*_ ≠ 0), and reversible strategies (characterized by bidirectional transitions *P*_*S*→*R*_ ≠ 0 and *P*_*R*→*S*_ ≠ 0). The “irreversible somatic differentiation” strategy corresponds to a specialized case of *RD*_*R*_ in our model, which we formally designate as 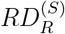. Similarly, reversible strategies represent a subset of *RD*_*RS*_, denoted as 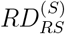. The superscript (*S*) explicitly indicates stage-independent differentiation probabilities [19].

Our analytical results confirm the previously predicted superiority of “irreversible somatic differentiation” 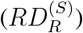 in large organisms, as illustrated in Fig. 4 and rigorously proven in Appendix A.3. Specifically, we establish that the reproduction rates of 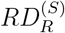 remain bounded in large organisms

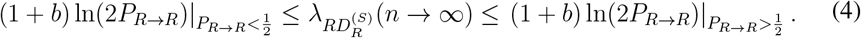

**Figure 4:**
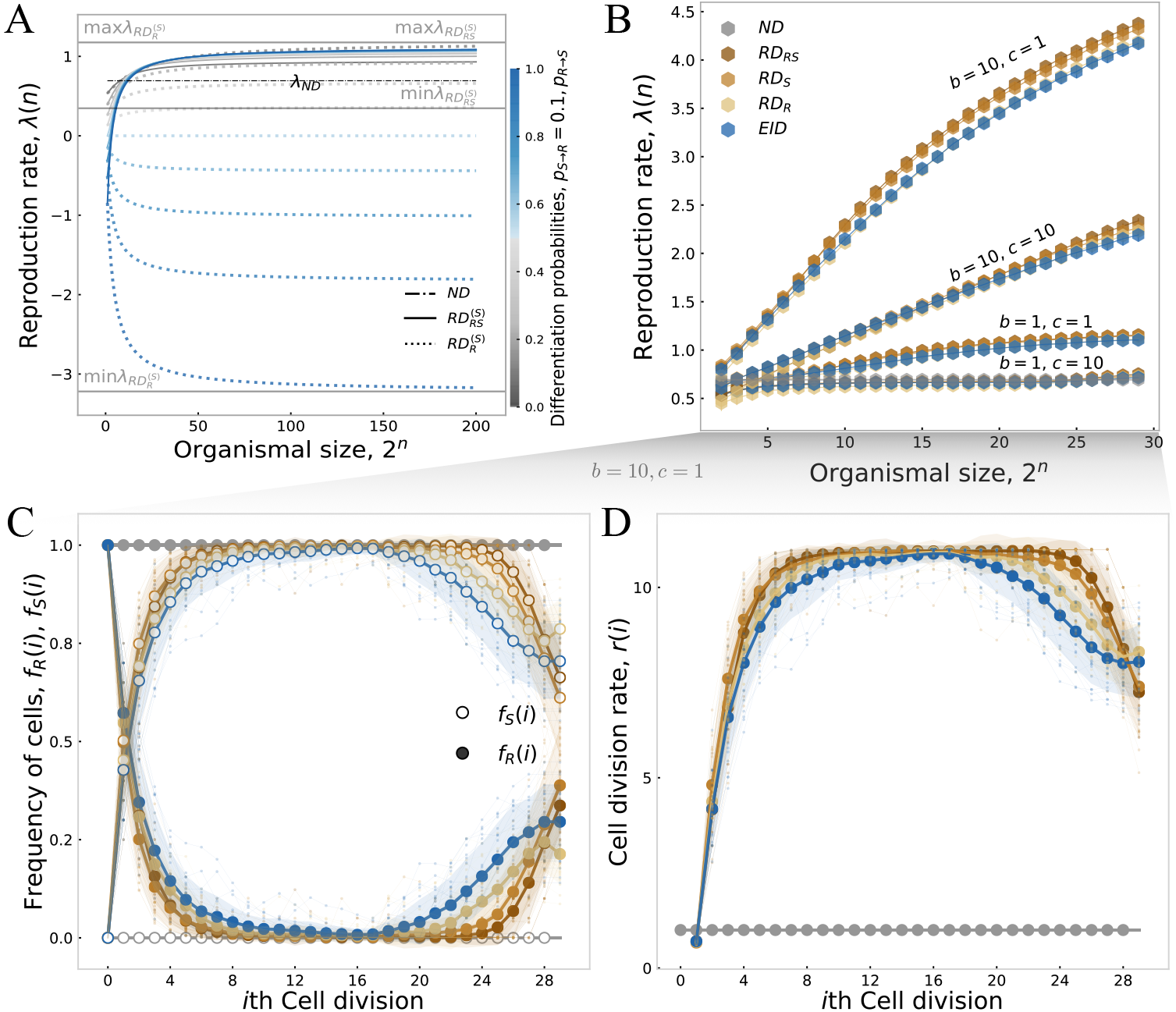
Comparison of optimal differentiation strategies under stage-independent and stage-dependent cell differentiation in large organisms. The reproduction rates of optimal stage-independent strategies (**A**) across cell division times were analyzed with fixed *P*_*S*→*R*_ = 0.1 and *P*_*R*→*S*_ varying from 0 to 1 in 0.1 increments. Analytical boundaries (gray lines) defined the reproduction rate limits (min *λ*, max *λ*) for each strategy category, with parameters *α* = *β* = *b* = *c* = 1. Stage-dependent strategies (**B**) exhibited mean reproduction rates across four differentiation benefit-cost configurations, calculated from 100 replicates per parameter combination (error bars: ± SD). For the optimal strategies in **B** (*b* = 10, *c* = 1, *n* = 29), the mean fractions of *R* and *S* cells (*f*_*R*_(*i*), *f*_*S*_(*i*)) showed divergent trajectories across divisions (**C**), with filled/unfilled circles denoting *R*/*S* cells (shaded areas: ± SD; 20 replicates). Corresponding cell division rates (*r*(*i*)) in **D** displayed time-dependent modulation, with shaded areas indicating variability. Numerical implementations followed the protocol in Appendix A.2 for *λ*(*n*) calculation.

Hereafter we denote *λ*(*n* → ∞) as *λ*(∞) for notational simplicity. By applying the limit *P*_*R*→*R*_ → 1 to Equation (4), we derive

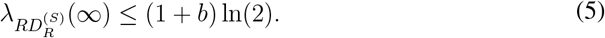

The reproductive rate bounds for 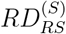 are established through

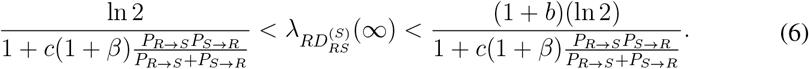

Through additional algebraic manipulation (see Appendix A.3), these bounds simplify to

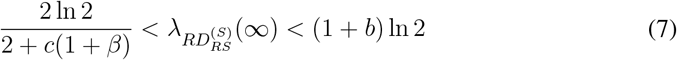

The comparative analysis of Equations (5) and (7) reveals that 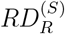 demonstrates competitive superiority over 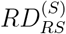 in organisms with large body sizes. Notably, the differentiation cost exhibits minimal influence on reproductive rates for stage-independent differentiation strategies in larger organisms.

While the constant reproduction rate of *ND* remains fixed at ln 2, both 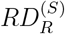 and 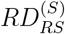 achieve selective advantage over *ND* under any beneficial conditions (*b >* 0). Notably, 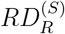 displays an intriguing mathematical property where reproduction rates can become negative. This phenomenon stems from the exponential decay term (2*P*_*R*→*R*_)^*n*^ in *R*-cell populations, which approaches zero as *n* → ∞ when *P*_*R*→*R*_ *<* 1, leading to negative rates through Equation (3). Although biologically unrealistic, these negative values theoretically correspond to sub-unity offspring production per individual per time unit. In contrast, 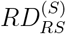 maintains stable *R/S* cell ratios through equilibrium dynamics, ensuring positive reproduction rates as *n* → ∞. We further prove that 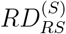 consistently maintains reproduction rates exceeding 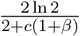, with detailed derivation provided in Appendix A.3. Analytical results in Appendix A.4 reveal fundamental differences in optimization trajectories: optimal 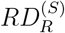 strategies approach *P*_*R*→*S*_ → 0, while optimal 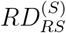 strategies evolve toward *P*_*S*→*R*_ → 0.

However, unlike stage-independent strategies, our analysis demonstrates that the *RD*_*RS*_ strategy achieves optimal performance in large organisms when cell differentiation probabilities exhibit state dependence (i.e., *δ*(*i*) ≠ 0), as shown in Fig. 4B. Notably, the reproduction rate differences remain minimal among cell differentiation strategies (all strategies except *ND*). This similarity arises from shared cellular differentiation trajectories across strategies. Specifically, these strategies may differ exclusively in their terminal cell division probabilities: *P*_*R*→*R*_(*n*) and *P*_*S*→*S*_(*n*). For an *EID* strategy with terminal probabilities *P*_*R*→*R*_(*n*) = 1 and *P*_*S*→*S*_(*n*) = 1, the *RD*_*RS*_ strategy would share identical cellular trajectories through the first (*n* − 1) divisions, deviating only through modified terminal probabilities *P*_*R*→*R*_(*n*) = 1 − *δ*(*n*) and *P*_*S*→*S*_(*n*) = 1 − *δ*(*n*), where *δ*(*n*) 1. Similarly, *RD*_*R*_ differs from *EID* only in *P*_*R*→*R*_(*n*), while *RD*_*S*_ diverges solely in *P*_*S*→*S*_(*n*). Consequently, all *RD* sub-strategies maintain reproduction rates comparable to the baseline *EID* strategy. The *RD*_*RS*_ strategy demonstrates distinctive advantages in large organisms through two synergistic mechanisms. First, increased organismal size enables sustained maintenance of *S*-cell fractions across multiple division states (plateau states in Fig. 4CD), thereby enhancing cell division rates. Second, large organisms employing *RD*_*RS*_ can generate substantial *R*-cell offspring due to the synchronous binary cell divisions despite reduced *R*-cell fractions. These advantages scale with organismal size, as increased division stages provide more opportunities for *RD*_*RS*_ to optimize cell switching probabilities and maximize reproduction rates.

## 4 Conclusion and discussion

This theoretical study examines how organismal size governs the evolution of extreme irreversible differentiation (*EID*), defined as differentiation probability sequences approaching zero for generating alternative cell types. By modeling evolutionary dynamics through an expected organismal reproduction rate (offspring per unit time), we identify size-dependent optimization principles for cellular differentiation strategies. Systematic maturity size variations reveal a nonlinear relationship: intermediate-sized organisms maximize *EID* promotion, whereas larger organisms evolutionarily suppress *EID* through modified differentiation trajectories. The evolutionarily optimal strategy combines non-differentiation (*ND*) and reversible differentiation (*RD*) features, with *ND* dominating in small organisms and *RD* prevailing in larger counterparts. Larger organismal size delays the developmental timing of cellular irreversibility across both cell lineages. This temporal delay enhances fitness through two mechanisms: (1) increased accumulation of differentiation benefits via amplified *S*-cell production, and (2) reduced differentiation costs through minimized differentiation probabilities. Crucially, we provide formal mathematical proof that 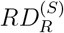 constitutes the optimal strategy for large organisms at *σ* = 0, resolving critical uncertainties from previous studies [19, 21]. These findings directly challenge conventional size-differentiation paradigms by demonstrating a reduced evolutionary preference for extreme irreversible differentiation in larger organisms compared to smaller ones, suggesting fundamental size constraints on cellular commitment evolution.

A major challenge in developmental biology involves classifying the complex differentiation trajectories observed during organismal development into distinct strategic patterns [35, 36]. While our model adopted the established methodology of categorizing differentiation strategies based on terminal division probabilities [21], this approach carries inherent limitations in distinguishing genuine differentiation strategies. To address this critical issue, we implemented a temporal analysis framework that tracks the developmental onset of differentiation irreversibility across cell lineages. This novel perspective reveals the precise developmental window when specific cell fates become irreversible, providing resolution for analyzing cellular decision-making dynamics. Our results demonstrate that *R*-type irreversibility emerges during early developmental stages under conditions of elevated differentiation costs, whereas *S*-type irreversibility initiates earlier when differentiation costs are reduced. This temporal decoupling of differentiation mechanisms suggests substantial plasticity in the developmental timing of terminal cell specification, even among phylogenetically related organisms with comparable body sizes. Furthermore, we identified a consistent pattern of delayed cellular irreversible differentiation in larger organisms, a finding that corroborates empirical observations of late germ-soma differentiation in perennial plant species [37]. Of particular significance is our discovery that reversible differentiation strategies become exclusively selected in organisms with indeterminate growth potential. This critical relationship between organismal size and differentiation commitment timing reveals an underappreciated constraint on developmental program evolution, suggesting that body size scaling laws may fundamentally shape the establishment of irreversible differentiation patterns across multicellularity.

While prior investigations into cell differentiation have extensively examined differentiation efficiency across organisms [18, 23, 24, 27, 29, 30, 38, 39], the relationship between organismal size and differentiation dynamics remains comparatively underexplored. This gap persists particularly in theoretical frameworks addressing the *origin* of cellular specialization, where most models assume a fixed number of organisms rather than organisms with growing cell numbers. Recent work by Cooper et al. provides critical insights, demonstrating an evolutionary preference for coordinated specialization in small cell groups where spatial proximity reduces coordination costs. These findings align with fundamental biophysical constraints, as smaller organisms inherently benefit from reduced intercellular communication distances that facilitate cooperative interactions. Parallel research by Yanni et al. reveals nuanced size-dependent effects, reporting enhanced differentiation in tree-structured cellular architectures but null effects in ring topologies with increased organismal size. However, their framework operationalizes differentiation through static task-allocation differences among predefined cell groups, fundamentally contrasting with our dynamic developmental perspective. Specifically, while existing approaches characterize fixed cell-type distributions in mature organisms, our model conceptualizes differentiation as an emergent property of growth-mediated cellular interactions. This distinction proves crucial when considering developmental trajectories, as our model captures the temporal progression of specialization patterns rather than analyzing static endpoints.

The interplay between organismal size and cellular differentiation has been frequently acknowledged in diverse biological contexts, yet systematic investigations into their causal relationships remain limited. For example, while Erten and Kokko [6] established that larger organisms tend to minimize differentiation probabilities under oncogenic risks, their framework employed static differentiation probabilities (mathematically equivalent to the *σ* = 0 condition in our system), where cell differentiation leads to cumulative effects of cellular damage. Our parameter analysis shows that when differentiation carries more costs – as implicitly presumed in their framework – evolutionary processes inevitably select strategies with less differentiation. The extreme situation is choosing non-differentiation (*ND*) strategies under high-cost conditions. Current literature exhibits notable gaps in addressing the size-dependent evolution of differentiation mechanisms. Previous comparative analyses of cellular specialization have involved the effects of organismal size, whereas it primarily focused on dichotomous comparisons between coordinated and stochastic processes [31]. Meanwhile, prior studies have explored either the impact of organism size or differentiation patterns [19, 21], however, our model establishes the first unified approach integrating organism size and differentiation trajectories via developmentally staged cell division patterns. This analysis reveals previously unrecognized interactions between organismal size and adaptive selection forces in guiding cellular fate determination.

Although our mathematical framework provides valuable insights for exploring irreversible cell differentiation dynamics across varying organism sizes, several limitations warrant consideration. The model’s foundational assumption that differentiation benefits exclusively depend on the relative proportions of *S* cells during organismal growth phases. While our selection parameters penalize mature organisms composed entirely of *S* cells, the optimal differentiation strategy permits transient homogeneous *S* cell configurations during developmental stages – a pattern particularly prominent in larger organisms. This finding contrasts with natural biological systems where differentiated cells frequently exhibit lateral inhibition mechanisms that actively prevent homogeneous cellular distributions [22]. Our idealized framework further assumes perfect cellular mixing and spontaneous cooperation, potentially overlooking the organizational challenges associated with increasing biological complexity. Empirical evidence suggests that larger organisms typically require sophisticated spatial organization and long-range communication systems [40], features absent in our current formulation. Previous investigations have highlighted the critical role of cellular communication architecture in differentiation patterning [28], though inconsistent findings regarding communication necessity exist in the literature [31, 41]. Additionally, our focus on total cell numbers assumes uniform cellular dimensions, despite emerging experimental evidence demonstrating significant size variation even among genetically identical cell types [42]. These simplifying assumptions, while necessary for model tractability, highlight important avenues for future research incorporating spatial structuring, cell-cell signaling dynamics, and cellular size relationships.

## A Appendix

### A.1 Reproduction rate

In the model, we treat both the number of cells and growth time as continuous, distilling the stage-dependent process down to two quantities for calculating reproduction rate: the expected offspring number of *R* cells 2^*n*^*f*_*R*_(*n*) and the amount of growth time for an organism to grow *t*(*n*). The reproduction rate *λ*(*n*) can be calculated by the following equation

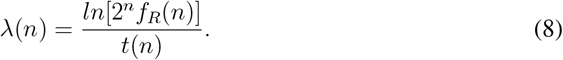

We use *f*_*R*_(*i*) and *f*_*S*_(*i*) to denote the fractions of *R* cell and *S* cell after *i*th cell divisions. Since each organism starts with a *R* cell, *f*_*R*_(0) = 1 and *f*_*S*_(0) = 0. *P*_*x*→*y*_(*i*) represents the transition probability from cell type *x* to *y* on *i*th cell division, where *x* and *y* are either *R* cells or *S* cells. After *i*th cell division, the expected fraction of *R* cells is *f*_*R*_(*i*) = *P*_*R*→*R*_(*i*)*f*_*R*_(*i* − 1) + *P*_*S*→*R*_(*i*)*f*_*S*_(*i* − 1) and the expected fraction for *S* cells is *f*_*S*_(*i*) = *P*_*R*→*S*_(*i*)*f*_*R*_(*i* − 1) + *S*_*S*→*S*_(*i*)*f*_*S*_(*i* − 1), which can be expressed in

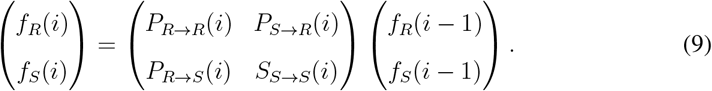

The expected *f*_*R*_(*n*) and *f*_*S*_(*n*) can be calculated recursively by equation (9)

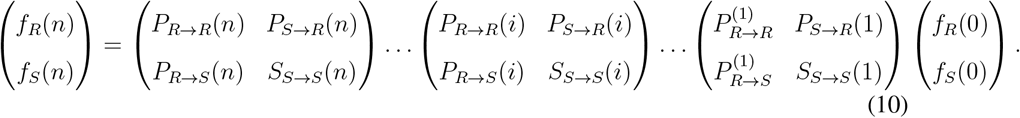

Since cells divide synchronously and no cell dies during growth, the expected *R* cells *N*_*g*_(*n*) and *S* cells *N*_*s*_(*n*) after *n*th cell division are

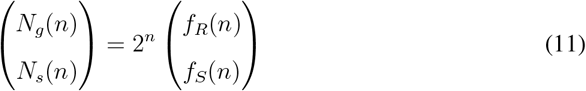

where 0 ≤ *f*_*R*_(*n*), *f*_*S*_(*n*) ≤ 1.

Cell division rate determines the growth time of organisms. Since cells divide with a rate 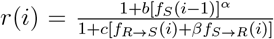 during ith cell division, the waiting time for a cell division *t*(*i*) follows the exponential distribution *f* (*t*(*i*)) = *r*(*i*)*e*^−*r*(*i*)*t*(*i*)^, where *f*_*R*→*S*_(*i*) = *f*_*R*_(*i* − 1)*P*_*S*→*R*_(*i*) and *f*_*S*→*R*_(*i*) = *f*_*S*_(*i* − 1)*P*_*S*→*R*_(*i*), see equation (9). Ths the expected waiting time from *i*th cell division to (*i* + 1)th cell division is 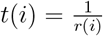. The expected growth time for organisms with total *n* cell divisions is

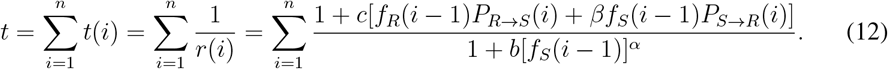

Substitute (11) and (12) into (8), we have

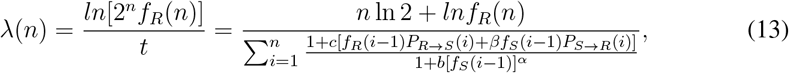

where *n* is the number of total cell divisions of organisms, *f*_*R*_(*i*) and *f*_*S*_(*i*) are fractions of *R* cell and *S* cell after *i*th cell division, *P*_*R*→*S*_(*i*) and *P*_*S*→*R*_(*i*) are the transition probabilities between *R* cell and *S* cell at *i*th cell division (1 ≤ *i* ≤ *n*). We have *f*_*R*_(0) = 1 and *f*_*S*_(0) = 0. For the non-differentiation strategy *ND*, no *S* cells are produced during growth, i.e. *P*_*R*→*R*_(*i*) = 1 and *P*_*R*→*S*_(*i*) = *P*_*S*→*R*_(*i*) = *P*_*S*→*S*_(*i*) = 0, *i* = 0, …, *n*. Therefore, *f*_*R*_(*i*) = 1, *f*_*S*_(*i*) = 0, *i* = 0, …, *n*. Thus from equation (13) the reproduction rate of *ND* which is denoted by *λ*_*ND*_(*n*) is ln 2. Biologically, the reproduction rate of *ND* means the number of cells doubles per unit of time.

### A.2 Numerical calculation of the reproduction rate

The calculation approach of the reproduction rate is the same as that in the previous model [21]. We use *R*_*RR*_(*i*) to show the probability of cell *R* produces two *R* cells at *i*th cell division. We first randomly choose a set of cell differentiation probabilities of cells at the first division step [*R*_*RR*_(1), *R*_*RS*_(1), *R*_*SS*_(1), *S*_*RR*_(1), *S*_*RS*_(1), *S*_*SS*_(1)]. We should note that [*S*_*RR*_(1), *S*_*RS*_(1), *S*_*SS*_(1)] does not play any role in the first cell division. For simplicity, we constrain the value of probabilities with one decimal, and *δ*(*i*) = ±0.1. Since *R*_*RR*_(1) + *R*_*RS*_(1) + *R*_*SS*_(1) = 1, thus there are totally 66 combinations for *R*_*RR*_(1), *R*_*RS*_(1), *R*_*SS*_(1). Thus, there are a total 4356 combinations for the set of the first cell differentiation probabilities. Then for each of the following cell divisions, the probabilities change due to *δ*(*i*). In order to find the optimal strategy at a given parameter point, we first chose *M* = 1000 randomly sets of cell probabilities for the first cell division from the 4356 strategies. Then for each chosen set of probabilities, we randomly chose *SAM* = 100 generated strategies by randomly choose *δ*. Thus, there are totally 10^5^ strategies being chosen. Then we calculate their reproduction rates and choose the strategy leading to the largest reproduction rate. To find the optimal strategy precisely, we compare the optimal strategy obtained above with the strategies generated by changing *δ*(*i*) on the strategy. The generated strategies include the one removing the first set of probabilities and compensating it with a last set of cell probabilities, or removing *d*_*n*_ by compensating with an *d*_0_. The aim of modifying the strategy is to find the globally optimal strategy in a local area. The seeking process will stop until it leading to the steady largest reproduction rate in its local neighbourhood.

### A.3 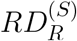 **is the only evolutionary outcome for larger organisms under** *δ*(*i*) = 0

When *δ*(*i*) = 0, cell differentiation strategy processes the same set of cell differentiation probabilities at different cell divisions during an organism’s growth, i.e. *δ*_*R*_(*i*) = *δ*_*S*_(*i*) = 0, where 1 ≤ *i* ≤ *n*. Thus the transition probability matrices (equation 9) of the cell distribution of an organism are the same for all cell divisions. For convenience, we denote the transition probability matrix without the cell division parameter 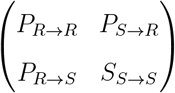.We can get the solution of cell distribution at each *n*th cell division [43],

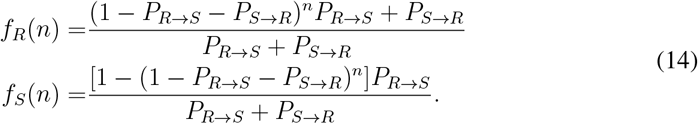

Substitute equation (14) into equation (13), we have the reproduction rate of stage-independent cell differentiation strategy

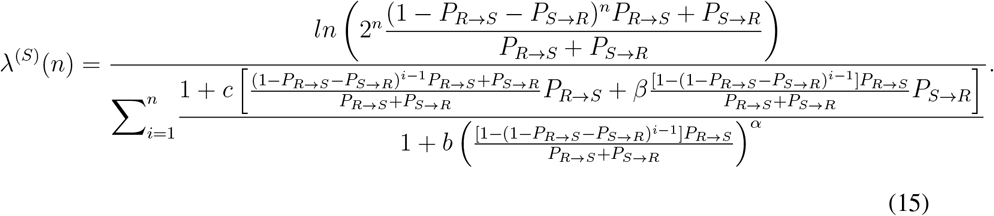

Consistent with previous results, our numerical results show the “irreversible somatic differentiation” 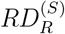 can evolves under intermediate cell differentiation benefits and costs, see Fig. 5A. This is due to the characteristics of cell differentiation rates and fractions of cell types of 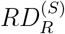 across cell divisions, which are eventually due to the features of cell differentiation probabilities of 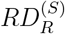, see Fig. 5A.

**Figure 5:**
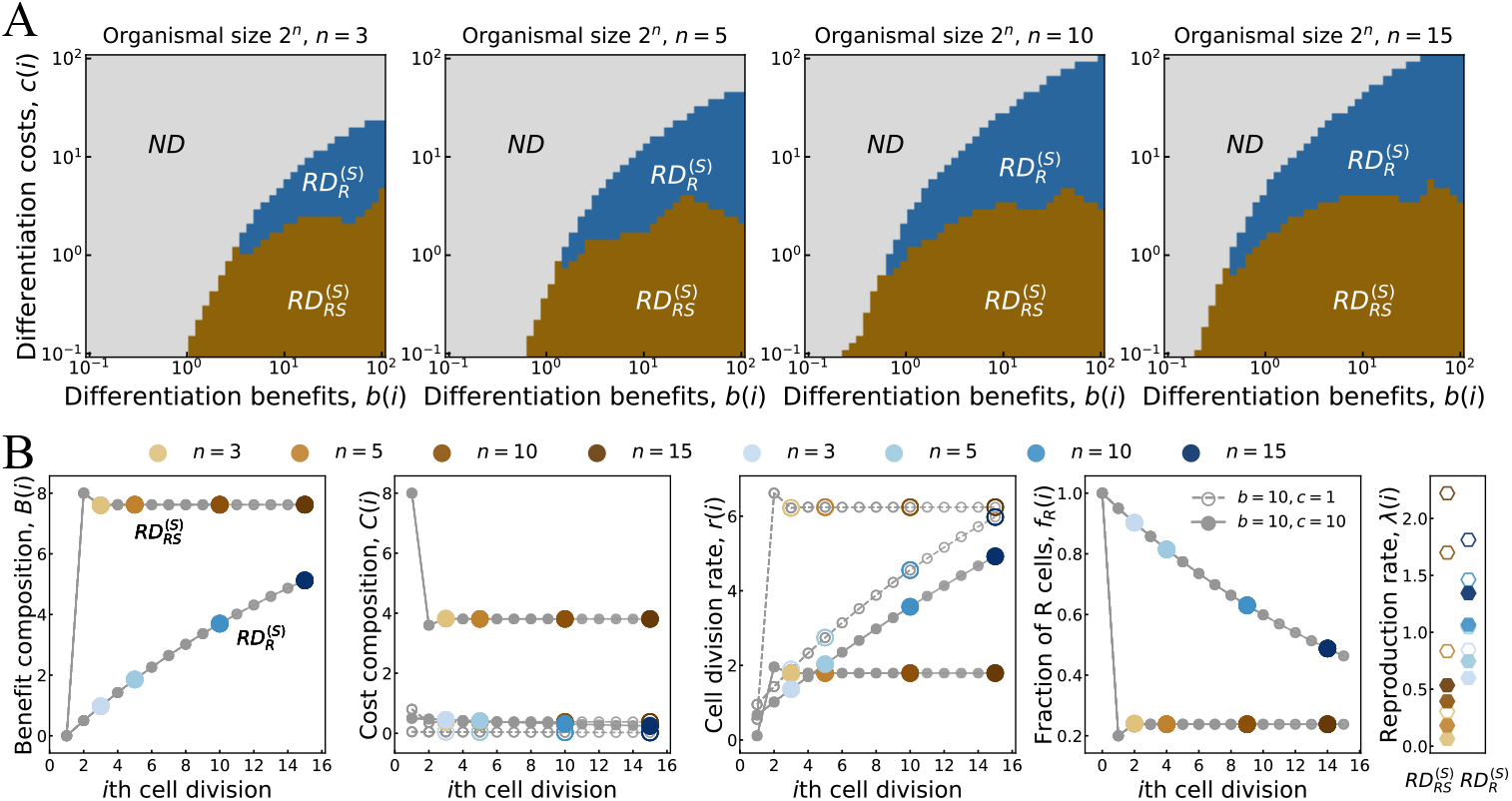
Organismal size changes the evolutionary likelihood of the optimal differentiation strategies under *δ* = 0. **A**. The distributions of optimal strategies under cell differentiation benefits *b* and costs *c* under maximal cell divisions *n* = 3, 5, 10, 15. **B**. The features of benefit composition *B*(*i*), cost composition *C*(*i*), cell division rate *r*(*i*), fraction of *R* cells *f*_*R*_(*i*) and reproduction rate *λ*(*n*) of the optimal 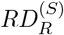 and 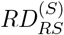 at *b* = 10, *c* = 1 and *b* = 10, *c* = 10 respectively. As *δ* = 0, thus different sized organisms share the same trajectories under the same optimal strategy, which are indicated by the grey dots. Solid circles and pentagons represent the feature values under *b* = 10, *c* = 10 and unfilled circles and pentagons represent that under *b* = 10, *c* = 1. We found that optimal strategies have smaller reproduction rate at high costs. Parameters of all panels: the maximum change of two successive differentiation probabilities −0.1 ≤ *δ* ≤ 0.1.

Together with equation (14) and 1 − *P*_*R*→*S*_ = *P*_*R*→*R*_, the reproduction rate of 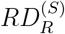 (*P*_*S*→*R*_ = 0) is

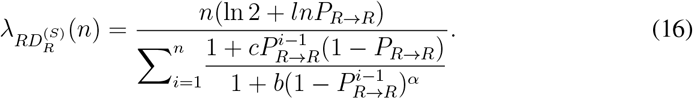

Next, we prove the bounds of 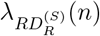. First, we look at a function *g*(*x*)

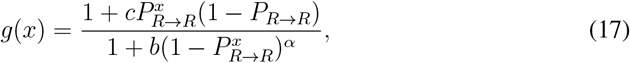

where *x* ∈ [0, ∞). The the derivative of *g*(*x*) is

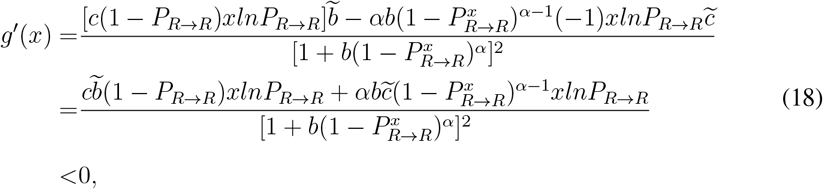

where we use 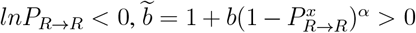 and 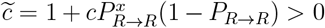 in the last step. Since *g*^t^(*x*) *<* 0, *g*(*x*) is a decreasing function in the interval [0, ∞). The maximum value of *g*(*x*) is *g*(0) = 1 + *c*(1 − *P*_*R*→*R*_) and the minima of 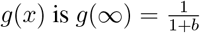.

We denote the denominator of equation (16) as 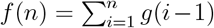. Thus *f* (*n*) is a discrete concave function, and

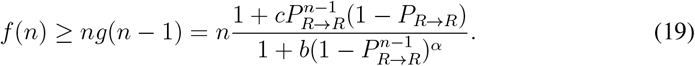

We next obtain the limits of reproduction rate 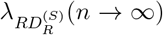 by inequality (19) and equation (16). Substitute inequality (19) into equation (16) and cancel the *n* in the denominator and numerator. Since the sign of 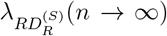 is determinate by the value of *P*_*R*→*R*_ as *lnP*_*R*→*R*_ *<* 0 when 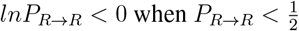 and *lnP*_*R*→*R*_ *>* 0 when 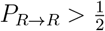. If 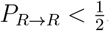, we have

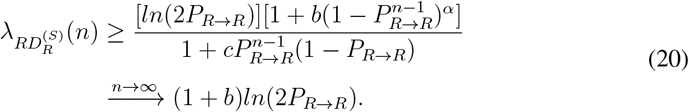

If 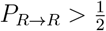, we have

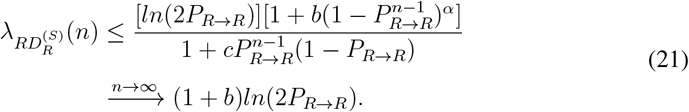

Together with inequality (20) and inequality (21), we have

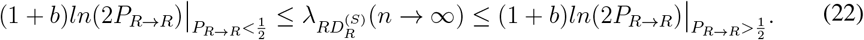

In reality, a negative reproduction rate is meaningless, thus, we are more interested in the case of 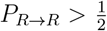. However, the theoretical conclusion of 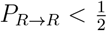 provides us the amplitude of 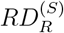. The above proof shows that only cell differentiation benefit *b* determines the amplitude of 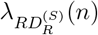. From (22), when 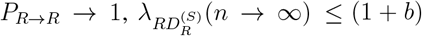 ln 2. The result indicates the reproduction rate of *RD*^(*S*)^ can be at most 1 + *b* times higher than *ND*. The reproduction rate of 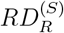 is confined by differentiation benefit *b*. The differentiation cost *c* is negligible.

For 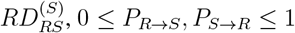, cell fractions will enter stationary state when *n* → ∞. We use 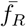 and 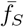 to denote the stationary distribution of the *R* cell and the *S* cell, thus

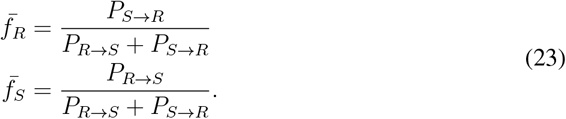

For any stage-independent *RD*, there exists an integer *M* which makes 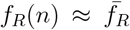 and 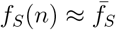 for *n > M*. By equation (15) and equation (23), we have

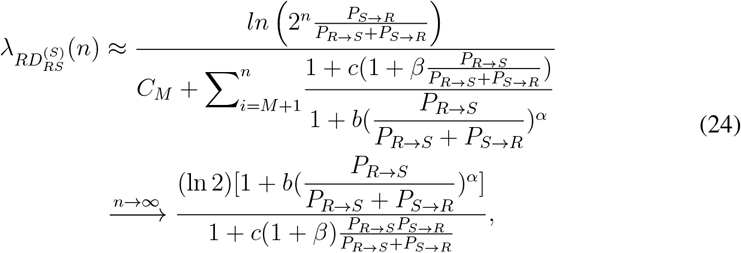

where 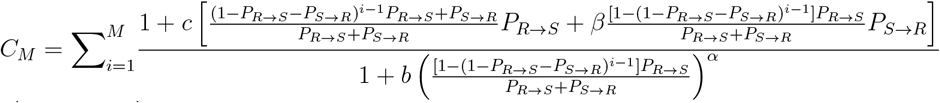 and 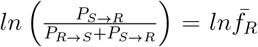 are constants and ignored. Since 0 *< P, P <* 1, thus 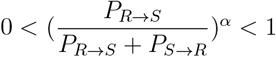. Thus *λ* 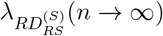 in equation (24) is bounded by

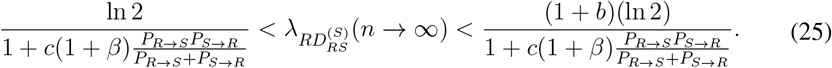

Furthermore, since 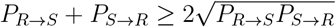 when *P*_*R*→*S*_ = *P*_*S*→*R*_, thus

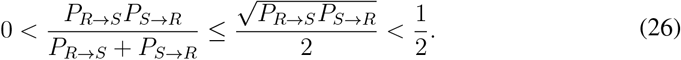

Substitute inequality (26) in to inequality (25), we have

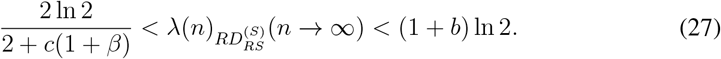

Inequality (22) and Inequality (27) show that 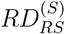 outcompetes other strategies in large organisms. Additionally, the cell differentiation cost *c* and the cost ratio of the *S* cell to *R* cell *β* together determine the minimum reproduction rate of 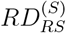. Unlike 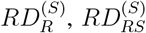 only has a positive reproduction rate when *n* is infinite.

#### A.4 Characteristic of cell differentiation probabilities of optimal strategy when *δ*(*i*) = 0 **and** *n* → ∞

From the main text, we know there are three strategies when 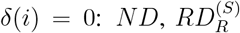 and 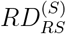. Next, we calculate the cell differentiation probabilities of the optimal strategy among 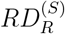 and 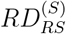. For convenience, we denote the reproduction rate of the each strategy when *n* is infinite as *λ*(∞), *P*_*R*→*S*_ as *p* and *P*_*S*→*R*_ as *q*. From equation (21), we have

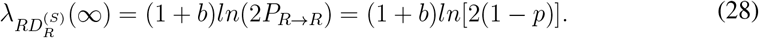

Thus, the optimal cell switching probability for 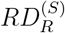 is *p* → 0 when *n* goes infinity.

Then from equation (24), we have the reproduction rate of stage-independent 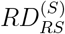 when *n* is infinity,

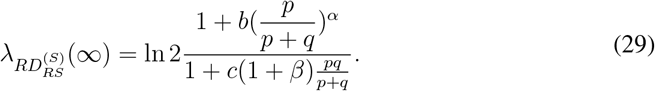

We next investigate the values of cell differentiation probabilities *p* and *q* under optimal 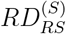 strategy when *n* goes infinity. 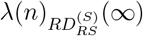 is a function with respect to *p* and *q*. The partial derivative of 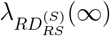 with respect to *p* is

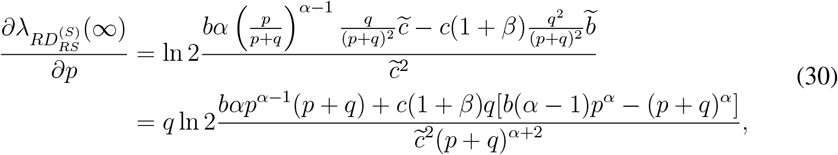

where 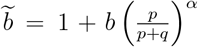 and 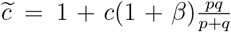. The denominator of equation (30) is positive. The first item of the numerator *bαp*^*α*−1^(*p* + *q*) is positive and the sign of the second item *c*(1 + *β*)*q*[*b*(*α* − 1)*p*^*α*^ − (*p* + *q*)^*α*^] is indefinite. With *α >* 1 and *B*(*i*) is large, 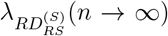 is a increasing function with respect of *p*. With *α <* 1 and *C*(*i*) are large, 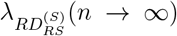 is a decreasing function with respect of *p*. The optimal cell switching probability of *p* depends on the value of *α, β, B*(*i*), *C*(*i*) and *q*. The partial derivative of 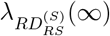 with respect to *q* is

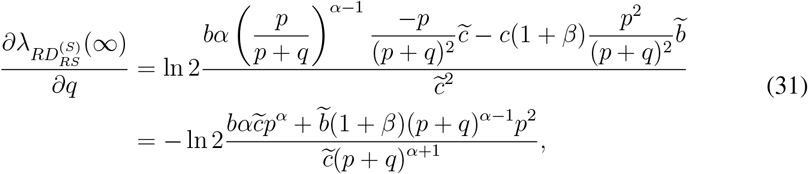

where 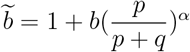 and 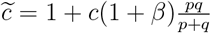. The numerator and the denominator are positive in the preceding equation. Thus, 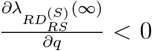. Thus, 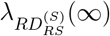 has the maxima value when the optimal cell differentiation *P*_*S*→*R*_ → 0.

### A.5 The fraction of the optimal cell differentiation strategies under varying organismal sizes

**Figure 6:**
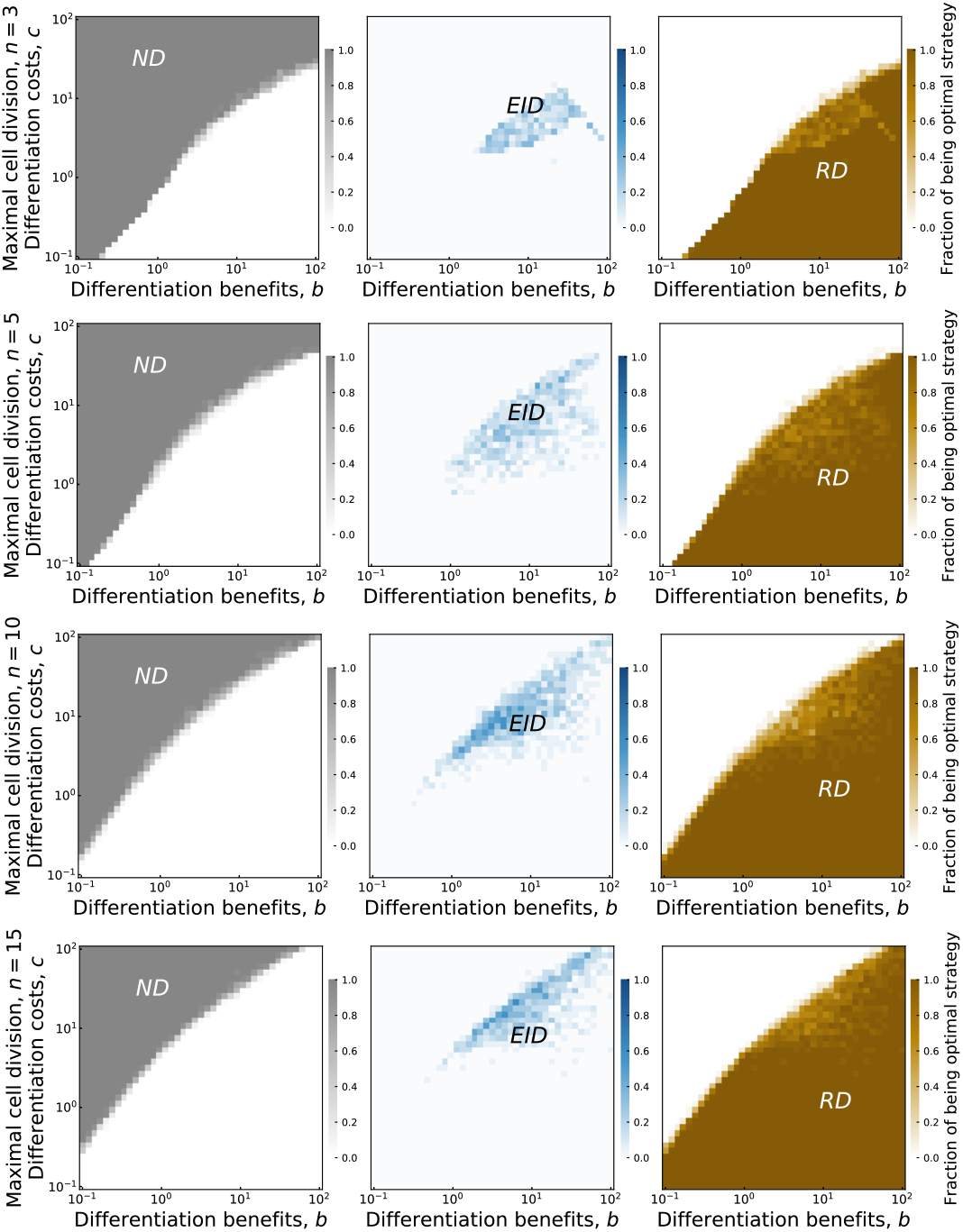
Intermediate organismal size promotes *EID*. The fraction of the optimal strategies are shown under the parameter space of cell differentiation benefit *b* and cost *b*. Maximum cell divisions are chose as *n* = 3, *n* = 5, *n* = 10 and *n* = 15. Parameters for all panels *α* = *β* = 1; the number of initial sampling *d*_1_, *M* = 1000; the number of stage-dependent strategies starting with a given *d*_1_, *SAM* = 100; duplicates at each pixel is 100, see appendix A.2 for more detail.

## Acknowledgments

*This work was supported by the National Natural Science Foundation of China [grant number 12401644] and the Natural Science Basic Research Program of Shaanxi Province of China [grant number 2024JC-YBQN-0005]*.

